# Acquisition of cell identity in the brown alga *Ectocarpus*: which of time, cell shape or position matters most?

**DOI:** 10.1101/2021.08.21.457218

**Authors:** Bernard Billoud, Denis Saint-Marcoux, Sabine Chenivesse, Carole Duchêne, Camille Noûs, Jane A. Langdale, Bénédicte Charrier

**Author notes:** Laboratoire BVpam UMR 5079, CNRS, Université de Lyon, UJM-Saint-Étienne, France. Laboratoire de Biologie du chloroplaste et perception de la lumière chez les micro-algues, UMR7141, CNRS, Sorbonne Université, Institut de Biologie Physico-Chimique, 75005 Paris, France.

## Abstract

During development, cells undergo simultaneous changes of different types that together depict cell “identity”. In the multicellular brown alga *Ectocarpus sp*., while ageing, cells change shape and relative position within the filament. Understanding how these factors act and interact to specify cell identity requires markers of cell identity and the ability to genetically separate age, shape and position. Here we used laser capture microdissection (LCM) to isolate specific cell types from young sporophytes of *Ectocarpus*, and performed differential RNA-seq analysis. Transcriptome profiles of cell types in the wild-type strain provided signatures of the five cell types that can be identified by shape and position. In two mutants, where the relationship between cell shape, position and age are altered, transcriptome signatures revealed that little differential expression could be identified when only shape was perturbed. More generally, although the two mutants are characterised by opposite morphological phenotypes, their transcriptomes were remarkably similar. We concluded that despite the robustness of cell differentiation during WT development, neither the shape nor the position of the cell could serve as a faithful gauge for tracking differentiation.

## Introduction

Morphogenesis in any organism requires that cells differentiate in precise spatial and temporal domains, often in the context of growth. Understanding how differentiation is regulated requires a consideration both of what defines the identity of any particular cell and of the processes that impart that identity. A cell is usually first defined by its shape, i.e. the way it looks under microscope. Next, with additional experimental tools, cell identity can be defined more precisely by the behaviour of the cell, e.g. its cell division rate or its metabolic activity. In this context, gene expression activity can be used as a snapshot of the cell identity state at any given time point or spatial position, either during normal growth or in response to stress. The total number of genes expressed per cell can then be used to assess the extent to which differentiation has progressed (Gulati et al., 2020), whereas the diversity of expressed genes (transcriptome signature) can be used as a proxy of cell identity (Chen et al., 2015). A wealth of data on hallmarks of cell identity in metazoan and plant cells has been acquired through single-cell transcriptomics (scSeq) (Zhang et al., 2021) (reviewed in plants in Seyfferth et al., 2021).

The acquisition of cell identity has traditionally been considered in terms of intrinsic (lineage) and extrinsic (positional) regulation. In plants, the external environment also plays a major role and many studies have reported ‘Omics’ data in a variety of plants subject to different environmental stresses (Zandalinas et al., 2021). Very few intrinsic signals triggering cell identity have been identified but cell shape has been widely shown to control cell fate in metazoans (Chen et al., 2020; Luxenburg and Zaidel-Bar, 2019). For example, long and short morphotypes of cardiomyocytes of similar age have transcriptome signatures which differ from the normal type. Both exhibit different numbers of total transcripts and a reduction in expression of signaling-related genes (Haftbaradaran Esfahani et al., 2020). Cell age is another intrinsic regulator, with cell maturation leading to modification of the structural organisation of the nucleus, in turn leading to gene expression and metabolic changes (Lans and Hoeijmakers, 2006; da Silva and Schumacher, 2021). Extrinsic regulators are more common, however, and cell position was also shown to control cell fate in many examples in both animals and plants. In the multicellular green alga *Ulva*, the position of cells at the 4-cell stage was also shown to be more important for cell differentiation in the thallus than an increase in genetic complexity (Fjeld and Løvlie, 1976). Such positional information can be imparted from neighbouring cells through biophysical or biochemical signals. For example, in plants where cells are constrained by a cell wall, growth and division of any individual cell requires co-ordinated adjustment in biophysical properties of neighbouring cells, as shown during the emergence of lateral roots (Vermeer et al., 2014). Many cellular differentiation processes in plants are additionally regulated by biochemical spatial information, for example provided by a polarised flux of the signaling phytohormone auxin, the perception and response to which depends on the position of a cell within a tissue, with differential activities seen after the first division of the zygote (Winnicki, 2020). In metazoans, cell position within a tissue also influences differentiation from an early developmental stage. For example, in the early embryos of mammals, the fate of the trophectoderm and inner cell mass (future embryo proper), hinges on the position of the cells – outer or inner – relative to the surface of the blastocyst (Mihajlović and Bruce, 2017). Notably, the tight inter-relationship between cell shape, cell age and cell position makes it difficult to identify primary regulators of cell identity in many organisms.

The shape (e.g. Zegman et al., 2015) and age (e.g. Kordyum et al., 2019; Zhang et al., 2005) of a given cell type can often be modified physically or genetically but altering cell position without introducing simultaneous changes of a biochemical or biophysical nature is less tangible, particularly in organisms with cell walls. In this case, organisms comprised of uniseriate filaments, where each cell has only two neighbours (hence limited positional input), and cell age and shape can be easily identified, are good models for distinguishing the effects of cell shape, age and position on cell differentiation. Strings of cyanobacteria, fungal hyphae, moss protonemata or algal filaments fulfill most of these criteria but not all. For example no cell shape changes occur in the moss *Physcomitrium* or in fungal hyphae, and in the cyanobacterium *Nostoc* cell differentiation is triggered by external signals (nitrogen starvation). We have therefore used the filamentous brown alga *Ectocarpus* to dissect the extent to which cell shape, age and position influence cellular differentiation. Using LCM and cell-type specific RNA-seq in wild-type and in mutants displaying abnormal combinations of age, shape and position, we show that transcriptome signatures of cell-type identity are lost in the mutants, and that differences between the WT and mutant transcriptome cannot be directly attributed to shape, age or position.

## Material & methods

### Wild type strains and cultivation

Two WT strains: Male Ec32 (CCAP 1310/4; origin San Juan de Marcona, Peru) and female Ec568 strain (accession CCAP 1310/334; origin Arica, Chile) were used to perform crosses with the mutants (see below). Thalli were grown in half-strength Provasoli-enriched (Starr and Zeikus, 1993), autoclaved natural seawater (pH 7.8) in Petri dishes located in a controlled environment cabinet at 13°C with a 14:10 light: dark cycle (light intensity 29 mmol photon·m^-2^·s^-1^) as described in Le Bail and Charrier (2013). Light intensity and duration, temperature, external mechanical forces and gravity were maintained constant during the experiments.

For LCM experiments, filaments were grown in PEN slides (ThermoFisher Scientific LCM0522), as described in Saint-Marcoux et al. (2015). Three slides were simultaneously cultivated in separate Petri dishes for each of the WT and the two mutant strains, and were considered as independent replicates. Fixation was performed when the filaments reached ∼ 50 cells. WT series #2 was fixed 5 days after WT series #1 and #3. Mutants were captured in an other experiment, several weeks after the WT.

### Production of mutants

The phenotype of the mutant *etoile* (CCAP 1310/337) was generated as indicated in Le Bail et al. (2011). In order to clear the genome from non causal mutations, *etoile* was crossed with the WT female strain (Ec568) from which a female descendant was selected and back-crossed with the WT male Ec32. A female [*etl*] descendant of this second cross was used in this study.

The mutant *knacki* was produced by UV-B irradiation of WT parthenogenetic gametes as reported in Le Bail and Charrier (2013). It was then crossed with the female WT Ec568 and then back-crossed with the original WT male Ec32. A [*kna*] female descendant was selected for this transcriptomic study.

### RNA extraction

*Ectocarpus* sporophytic filaments were grown directly on PEN slides from spore germination and grown in standard culture conditions for 10 days. PEN slides with ∼ 30-cell filaments were fixed and tissue extracted by laser capture microdissection as described in Saint-Marcoux et al. (2015). RNAs were extracted and amplified as described in Saint-Marcoux et al. (2015). Each cell type (A, E, I, R, and B) was represented by 3 independent replicates (series 1, 2 and 3 for A1, A2, A3, etc) of about 200 cells captured from 3 different PEN slides grown simultaneously. While a single A cell per filament was dissected, groups of 2-4 adjacent cells of the other cell types were extracted from the same filament. Mutant cell types followed the same experimental procedures as the WT, but mutants and WT cultures were not grown simultaneously. Because of their phenotype, mutant cell types were defined based on their position along the filament only. Therefore, in both mutants, A stands for one apical cell, S comprises 2-3 sub-apical cells and C represents the central part of the filament (∼ 5 cells).

### RNA-seq analysis

For each sample, 10 Millions of Paired-End reads were produced with the HiSeq NGS technology (BGI, see Suppl Table 1). The RNA-seq reads were filtered using Trimmomatic (Bolger et al., 2014) and rRNAs were removed by SortMeRNA (Kopylova et al., 2012). The remaining reads were mapped onto the *Ectocarpus* genome and transcriptome V2 representing 18271 genes (Cormier et al., 2017) available in Orcae (Sterck et al., 2012) using bowtie (Langmead et al., 2009). Differential Gene Expression was detected using edgeR (McCarthy et al., 2012; Robinson et al., 2010) setting the maximum False Discovery Rate (FDR) to 5 × 10^−2^. The reliability of the NGS results was checked by Q-RT-PCR amplification of a series of genes found to be significantly differentially expressed between the A and R cell types (Suppl data). Biases in GO-term representation between samples were identified in the Gene Ontology Database (Ashburner et al., 2000; Gene Ontology Consortium, 2021) using R 4.1 (R Core Team, 2021) to perform a Fisher exact test and adjust p-values for multiple tests by the hommel method. For Principal Component Analyses (PCA), we restricted the analysis to nuclear genes having no sex-specific expression, as mutant strains are female. In a first run, we included all 17106 genes in a global analysis. Then, for each of these genes, we computed a Variance Ratio (VR) as the variance between the mean expression level (in TPM) in A, E, and R samples in WT, divided by the mean variance of expression levels between the three replicates of the same sample. The 2000 genes having the highest VR were used to find the main PCA axes for WT expression levels. Then, the gene expression levels in all five WT cell types, or mutant samples, were plotted onto the same graph using the previously defined main axes. When comparing different genotypes, only genes expressed in both were considered.

## Results

### Five cell types, each with a distinct shape, position and age coordinates, differentiate in WT Ectocarpus

The brown alga *Ectocarpus sp*. is an excellent model to study cell differentiation. In its sporophytic stage (Fig 1A&B), cell differentiation proceeds along a single axis in the context of a uniseriate filament (Fig 1C). Soon after germination, the zygote grows through the elongation and division of the very polarised apical (“A”) cell. Elongated (“E”), sub-apical cells result from successive axial divisions of the A cell. As the filament grows, the E cells progressively get rounder after they divide. Going first through an intermediate (“I”) stage, the cell eventually becomes fully round (“R”). Monitoring cell differentiation over 7 days in bright field microscopy showed that this process occurs in ∼ 4 days (Suppl Movie 1). Measurement of the cellular dimensions showed that R cells had a 44% increase in equatorial diameter and a ∼ 39% decrease in length relative to E cells, which resulted in a volume increase of ∼ 27%. Cells on which branches emerge are named “B” cells (note that branches never emerge on A cells).

**Figure 1:**
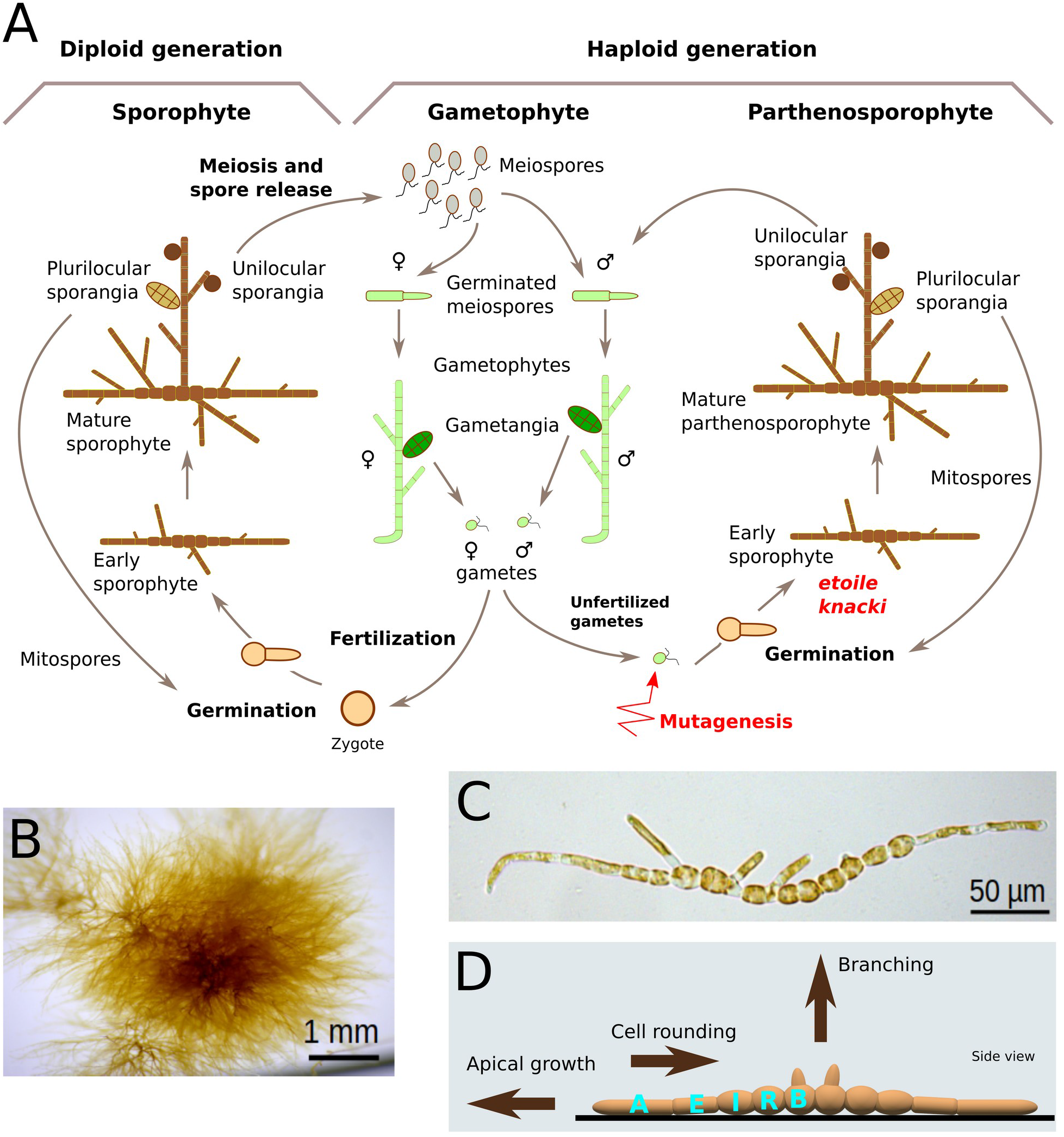
*Ectocarpus sp*. sporophyte filaments. display 5 cell types based on their shape, location and age. (A) Life cycle of *Ectocarpus sp*. The life cycle alternates between a haploid gametophytic stage and a diploid sporophytic stage. Both phases are filamentous. Unfertilised gametes germinate into parthenosporophytes resembling diploid sporophytes. Mutants *etoile* and *knacki* were produced by UV irradiation of unfertilised gametes (red arrow). (B) One month-old *Ectocarpus* sporophyte comprised of branched filaments. (C) 10-day old *Ectocarpus* sporophyte filament showing cylindrical cells at the distal end and spherical cells in the more central region, together with primary branches. D) Schematised representation of an early sporophytic filament. Three main cellular processes take place during growth. 1) apical growth, 2) cell rounding and 3) branching. Five cell types are defined along the filament, according to their shape, position and age.

As cells differentiate from A to R, filaments progressively elongate due to apical cell growth and division, so that R and B cells eventually reside in the most central portion of the filament (Fig 1D). Simultaneously, cells get older. In summary, cells at the tip of the filaments are the youngest and the most polarised because the filament grows by apical cell elongation and cell division, while cells in the centre are the oldest and the most isotropic shapewise because they result from progressive rounding. Therefore, in WT *Ectocarpus*, age, shape and position are intrinsically linked.

### Disentanglement of cell shape, age and position in two mutants

To disentangle the relationship between cell shape, cell location and cell age, we used two cell shape mutants, *etoile* and *knacki* (Fig 2). We divided the values observed for each parameter into categories, encoded by numbers (Fig. 2A&B). Shape is expressed as the the width/length ratio. Age is the (rounded) number of days since the last apical division. Position is an arbitrary numbering of domains within the filament, named A for the apical domain, S for the sub-apical, and C for the central domain.

**Figure 2:**
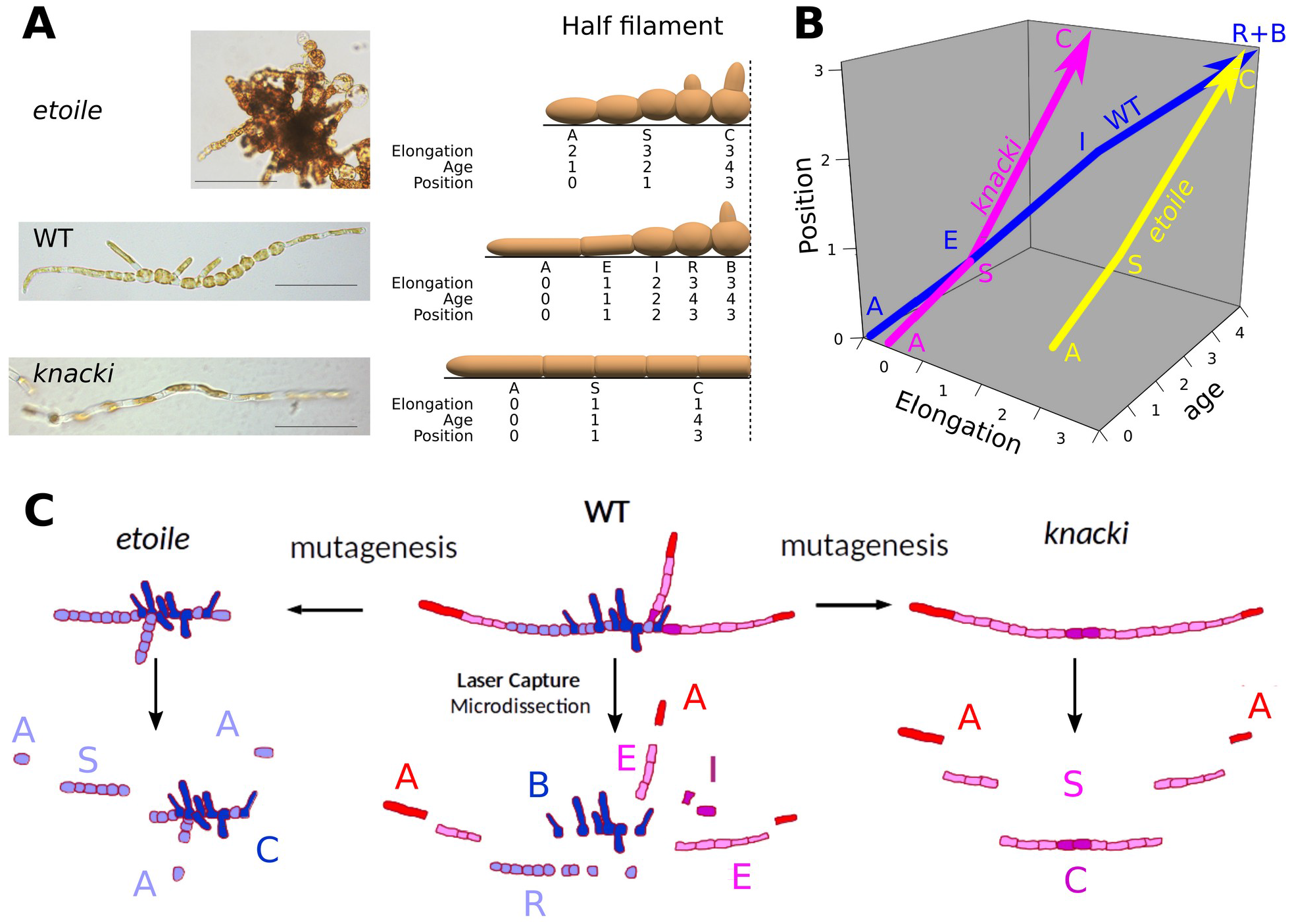
The impaired cell differentiation in the *Ectocarpus* mutants *etoile* and *knacki* allows to disentangle cell shape from the location within the filament and the cell age as second parameters. (A) Overall morphology of the mutants *étoile* and *knacki*, as compared to the WT. Photos of ∼ 2-week old organisms are shown, together with schematised half filaments. Note that in the WT, cell types are displayed by only one cell of each, but they are usually represented by several cells always grouped together. Scale bar = 50µm. Cell types for the WT and positional types for each mutant are characterised by (I) Elongation computed as the (discretised) length/width ratio, (ii) Age since “birth” of the cell issued from division of the apical cell, and (iii) Position relative to the global filament composition. To allow for categorial comparisons, all these parameters are discretised as integral values. (B) The three features of cell differentiation, shown as three spatial dimensions. Changes in cell state are represented as trajectories in this space, for WT (blue) and mutants *etoile* (yellow) and *knacki* (pink). (C) Laser ablation method, for sporophyte filaments of WT (center), and mutants *etoile* (left) and *knacki* (right). Each WT cell type is represented with a different color. Red for A (apical) cell type, pink for E (elongated) – both A and E are cylindrical – purple for I (intermediate stage in which cell shift from cylindrical to spherical cell shape), light blue for R (spherical cells) and dark blue for B (branched cells, initiating lateral apical growth). Mutants *etoile* and *knacki* are depleted in specific cell types: A (apical), S (sub-apical) and C (central), based on their position.

The first mutant, *étoile (etl)*, is characterised by an impairment in cell shape and apical growth: all cells appear bulky, and growth is slowed down, eventually stopping after about one week (Le Bail et al., 2011). As branching is initiated at the same pace as in the WT (Nehr et al., 2011), *etl* adult morphology looks bushy. More precisely, cells of both A and S domains are rounder than the WT A and E cells which respectively occupy the same domains (Fig. 2A). Their elongation state is shifted by two steps, and appears similar to the WT I and R cell types. Cells of the C domain have a round shape and overlap in term of position with that of WT R cells. This phenotype thus allows cell shape to be disentangled from cell position within the filament. In addition, since sub-apical cells are produced by slow apical cell divisions, cells from the A and S domains in *etl* are of similar ages as the WT E and I cells, respectively. Therefore, the early stages of *etl* allow cell age to be disentangled from cell position and shape (Fig 2B). The second mutant, *knacki* (*kna*), displays a phenotype opposite to *etl*. Round central cells are absent in *kna*, and branching is delayed, making the overall morphology well spaced-out (Fig 2A). Cell position and shape are similar to the WT for the apical cell. The shape of *kna* S cells is similar to that of WT E cells of comparable age and position. However, C cells fail to become round over time due to their more internal position within the filament (Fig 2B). Therefore, the later stage of *kna* development allow cell shape to be disentangled from cell age and cell position. Both mutants display unique parameter combinations that are not available in the WT.

### The five WT cell types have different transcriptomic signatures

We used laser capture microdissection (LCM) to manually isolate the five different cell types in WT filaments based on their morphological differences. A, E, I, R and B cell types were dissected from 2 week-old WT filaments (at ∼ 50-cell stage, Fig 2C). From the extracted RNA, 30 millions reads were obtained for each cell type, summed from 3 independent biological replicates (∼ 10^7^ reads each; see Suppl. Results, along with Figures and Table cited therein). Prior to analysing the expression pattern in different spatial domains along the filaments, we assessed the extent to which the transcriptome, as we captured it, was a reliable representation of the total *Ectocarpus* sporophyte transcriptome. In *Ectocarpus*, the sporophytic phase lasts as long as the gametophytic phase, i.e. about 6 weeks (Fig 1A) (Charrier et al., 2008). Interestingly, while 10-day old sporophytic filaments correspond to ∼ one ninth of the total life cycle, 97.5 % (18017 transcripts) of the total predicted transcriptome (18479) was expressed in these filaments. As such, only 462 genes are absent from our dataset. When comparing our data with Lipinska et al. (2019), we found that ∼ 80 % of the sporophytic-biased and ∼ 50 % sporophytic-specific genes were present in our dataset (Suppl Table 3). Correspondingly, (and more surprisingly) 45% of the gametophyte-biased and ≥ 10 % of the gametophyte-specific genes were also expressed in the early sporophyte. Altogether, our dataset contains as many sporophytic as gametophytic genes as identified by Lipinska et al. (2019) (about 1700 in each case), and represents over 97% of the predicted transcriptome.

Examination of transcriptomes in specific cell-types revealed that each of the five expressed about 77% of the total number of the genes, meaning that a significant proportion of the transcriptome is common to all cell types. However, comparison of transcript abundance revealed differences between cell-types. For example, 1134 genes were significantly differentially expressed (DE) between the A and the R cell types. Narrowing the analysis down to adjacent cell types, the comparison showed a progressive shift from the apical A cell to the central R cell. (Table 1). Indeed, 253 genes were DE between A and E, and 198 between E and I but only ∼ 40 genes were DE between the I-R and the R-B cell types. Therefore, the transition from A → E and E → I is marked by bigger transcriptional changes that the I → R, I → B and R → B transitions. This suggests that after a major reconfiguration between A and E cell types and the initiation of rounding from E to I, the I, R and B cell types represent a relative steady state in which gene expression becomes progressively stabilised (Fig 3A). Significantly, among the DE genes, those involved in photosynthesis and carbon metabolism were over-expressed in round/central/older cells, while elongated/peripheral/younger cells expressed more stress-related genes (Suppl File 1). Table 1 shows that the A cell type is more similar to the B cell type than to E, I and R, an observation that is easily explained because the B cells are initiating a new A cell through branching, and thus they combine I/R cell identities with the A identity. Collectively, these transcriptomes reflect the dynamics of cell differentiation in time and space, providing a spatial resolution as narrow as one cell (∼ 15 µm) and a temporal window of ∼ 1 day, and show that gradual changes in cell identity, as defined by shape, position and age, can be monitored by progressive modifications of transcriptome signatures.

**Table 1:** Number of genes showing differential expression (FDR < 5.10^−2^) in all pairs of cell types.

| Cell type | A | E | I | R | B |
| --- | --- | --- | --- | --- | --- |
| A |  |  |  |  |  |
| E | 253 |  |  |  |  |
| I | 870 | 198 |  |  |  |
| R | 1134 | 366 | 48 |  |  |
| B | 619 | 127 | 22 | 36 |  |

**Figure 3:**
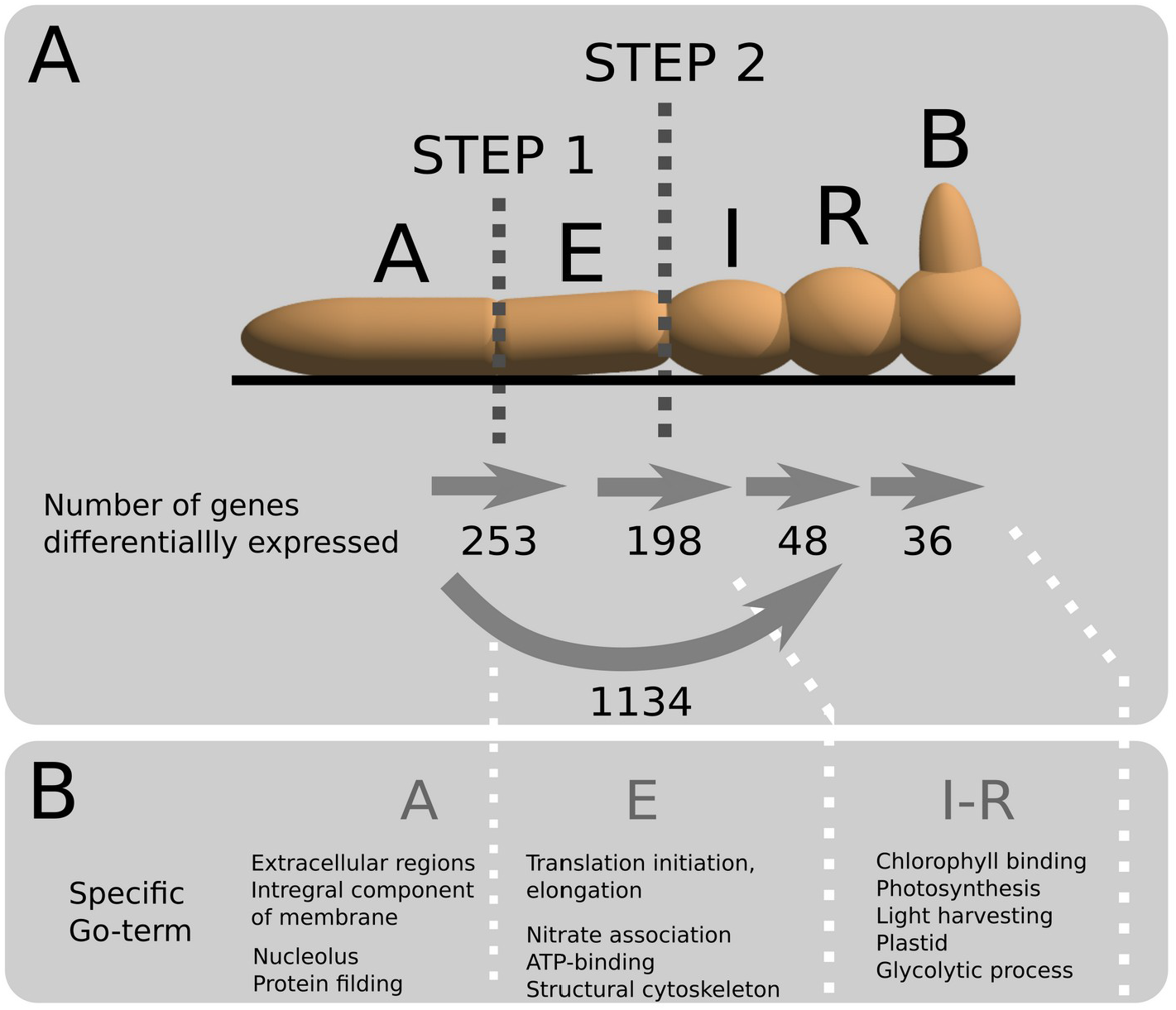
Gene expression is tightly regulated along the sporophytic filament of Ectocarpus. (A) Number of genes differentially expressed (DE) along the filaments. Two major steps occur for the transition A→E and E→I. Once cylindrical E cell starts rounding, expression profiles has stabilised and only few genes are differentially expressed. (B) GO-term significantly representative of the 3 domains A, E and I-R. For B cell type, see text.

To get insight into the specific function of the 5 different cell types, we analysed the GO-terms for genes that were significantly under or over-represented when comparing cell types in pairs (Fig 4; Suppl Table 5). We found that the GO-terms over-represented in the A cell type (negative values for X-A comparisons in Fig 4) are related to extracellular plasma membrane components, suggesting a function in the perception of external signals at the surface of the cell. The second and third GO-terms relate to the nucleolus and protein folding. These two functions are most likely linked, as the nucleolus is known to store and repair misfolded proteins in response to stress in animal (Frottin et al., 2019) and plant cells (Kalinina et al., 2018). Compared to A cells, E cells are involved in light harvesting and glycolytic process, suggesting that they are more active photosynthetically than A cells, yet to a lesser extent than the I and R cells. It appears that photosynthetic activity progressively increases while cells undergo transformation from A to R. Interestingly, E cells are also clearly marked by GO-terms specific of translational activities when compared with I and R cells. Less represented significant GO-terms of E cells are nitrate assimilation, cytoskeleton structural component and ATP-binding. GT-term analysis confirmed the intermediated state of B cells as reported in Table 1. and revealed that the main difference between A and B relates to activity in photosynthesis and translation (Fig 4). No differential GO-term representation could be found between I and R cell types, suggesting that they have similar identities. Collectively, the GO-term analysis, summarised in Fig 3B thus extends our understanding of cell differentiation as initially depicted in Fig 3A and reinforces the idea that differentiation proceeds with two main steps along the filament of *Ectocarpus*, rather than being linear.

**Figure 4:**
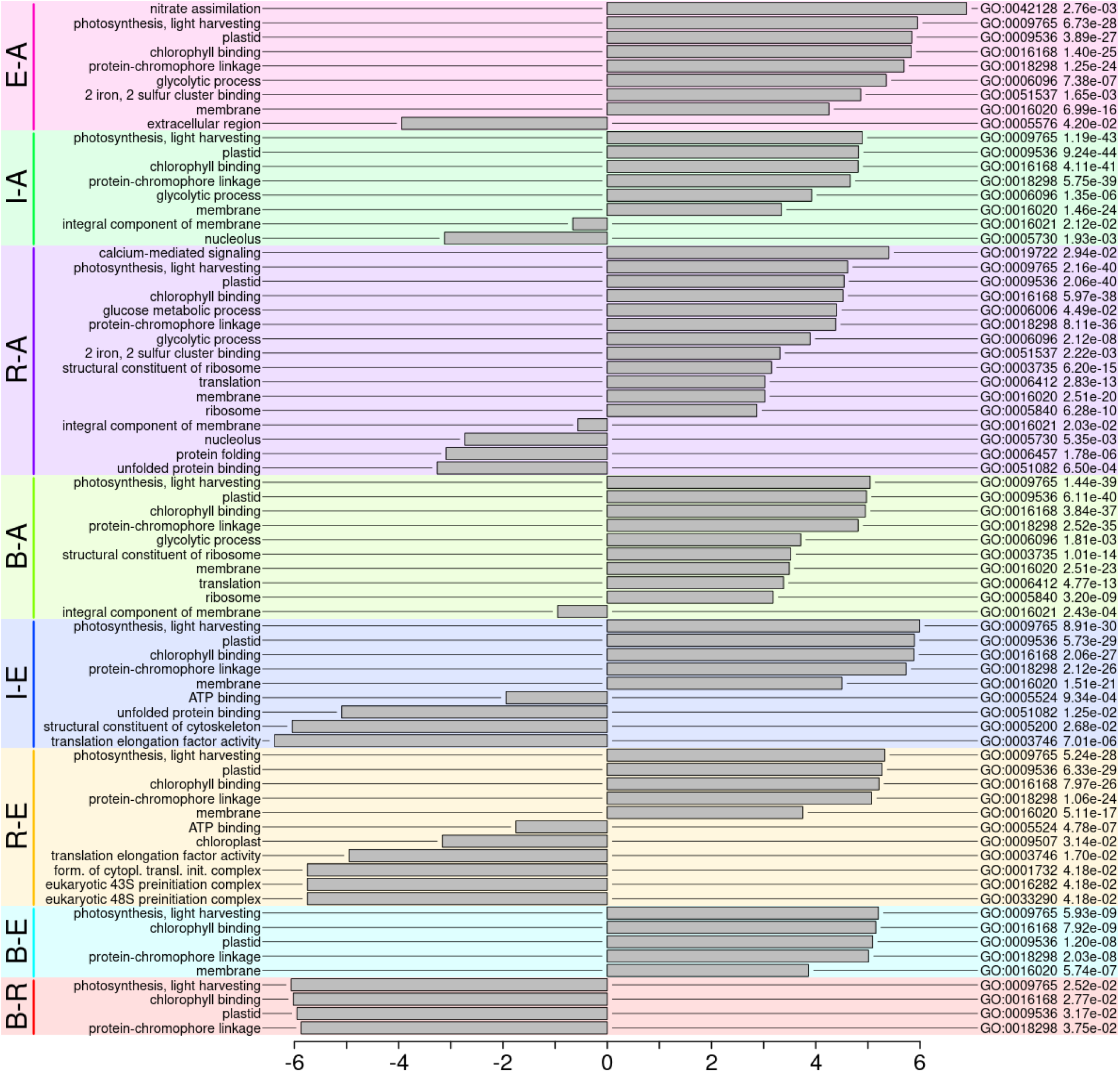
GO-term analysis displays specific functions for mainly 3 cell type domains. Differentially expressed genes are shown for pairs of cell types. The horizontal columns indicate the log2(fold change) of representation between two cell types of the pair. For a pair denoted as “X-Y”, a positive value for a given GO-term means that this term is over-represented in X as compared to Y. GO-term numbers and adjusted p-values are shown in the right-hand side columns.

### Transcriptome signature of cell type domains are lost in cell shape mutants

To determine the extent to which cell shape, as opposed to cell age or position in the filament, influences cell-type specific transcriptome signatures, we examined transcriptome profiles in *etl* and *kna* mutants. Because both *etl* and *kna* are cell shape mutants, the filament domains to be captured were defined only from the position of the cells within the filaments (Fig 2C). Hence, RNAs from apical (A), sub-apical (S) and central (C) domains were extracted, amplified and sequenced. Sequencing results were first analysed within each mutant, by looking at the number of genes that were differentially expressed in mutant cell types A, S and C. Within *etl* only 4 genes were differentially expressed in the different spatial domains of the filament, even with a very permissive FDR of 5×10^−2^ (Fig 5A). This suggests that all cell types are similar at the transcriptional level and that all the differences displayed between WT cell types are abolished in this mutant. Similarly, very few genes were shown to be differentially expressed between cell types in *kna* mutants, with just a single gene being DE between the apical A and sub-apical S domains (Fig 5B). This abolishment of DE between cell-types in two mutants that have different cell shapes but are united by having a phenotype in which cell shape is fairly consistent regardless of cell age or position within the filament suggests that cell-type transcriptome signatures are defined primarily by cell shape and not by age or position.

**Figure 5:**
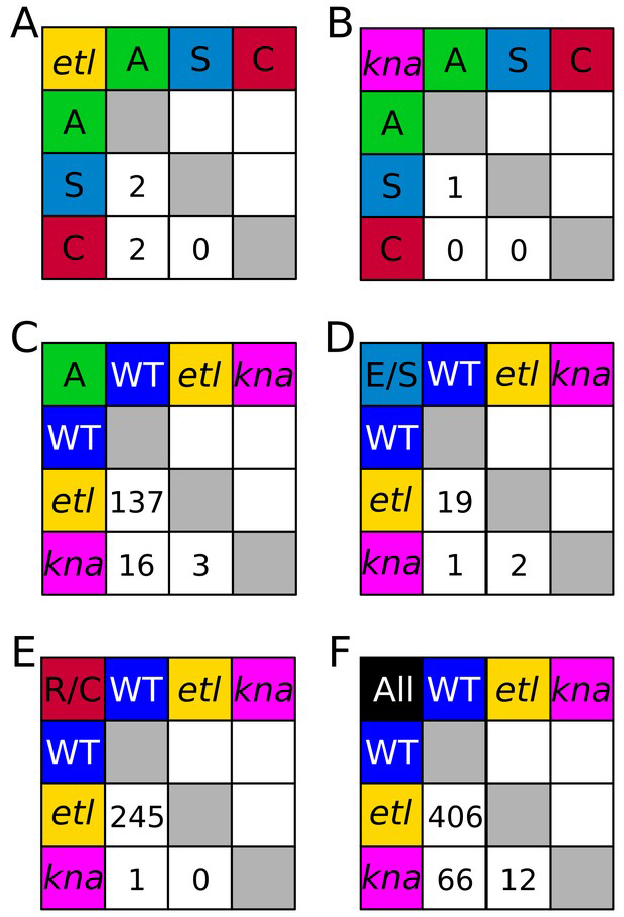
Number of genes differentially expressed (FDR < 5.10^−2^) between cell types or genotypes. (A) between spatial domains in *etoile*. (B) between spatial domain in *knacki*. (C) Between WT and mutants in domain ‘A’ (apical). (D) Between WT ‘E’ and mutants in domain ‘S’ (sub-apical). (E) Between WT ‘R’ and mutants in domain ‘C’ (central). (F) Between WT and mutants for all cell types and domains together.

In a second step, we assessed the impact of cell position on cell-type transcriptome signatures by comparing the mutant dataset with that of WT. 10446 genes were found to be expressed in common between the WT and *etl*, with similar representativeness in 3 replicates of the 3 spatial domains (Suppl Fig 7A). Between WT and *kna*, we found a lower number of common genes (7712) but comparable abundance (Suppl Fig 7B). For each spatial domain, we counted the number of genes that were DE between the WT and the mutants. First we examined cells in the apical position. Only 16 DE genes were observed between WT and *kna* apical cells, which are very similar in shape, while 137 genes differ between WT and *etl* apical cells, which do differ in shape (Fig 2A&B, Fig 5C). As such, cell shape not position is defining the transcriptome signature of the apical cell. However, the A spatial domain was found to be more similar between the two mutants than between any of the two mutants and the WT despite their opposed shapes.

Comparing WT profiles to the mutant profile in the S spatial domain (Fig 5D), however, revealed a different relationship between age, shape and position to that seen in the apical cell. Specifically, *kna* differs from the WT with respect to DE expression of 1 gene, consistent with the fact that sub-apical *kna* cells are similar in shape to WT E cells. But only 19 genes were found DE between *etl* and the WT, and 2 DE genes between the two mutants, although in both cases, elongation and age are different. Even more puzzling, when we compared the WT R to the C spatial domain (Fig 5E), we found *etl vs* WT displayed the biggest number of DE genes of all pairwise comparisons (245), while these cells have similar shape age and position. Conversely, *kna* had few, if any DE with the two other strains, despite the very special shape of its central cells (Fig 2A&B). When genes were pooled regardless of their type or spatial origin, *etl* and *kna* looked similar at the transcriptional level (only 12 DE genes) (Fig 5F). This surprising resemblance suggests that in *Ectocarpus*, the location of cells along the filament is not paramount in the control of gene expression. The most striking trend is that, although their phenotypes are opposed with respect to the WT, the two mutants differ by few DE genes.

To go further in the analysis, we performed systematic comparisons between pairs of types or domains for which exactly two parameters had different values. The results show that the largest differences are always found when comparing *etl* A with any WT cell type (Suppl Table 5). In all cases, the two mutants display few differences between them. This was confirmed by a Principal Component Analysis (PCA), for which, as the WT is male and the two mutants are female, sex genes were removed, leaving 17106 nuclear genes. By separating all the samples on the basis of the expression pattern of these genes, the main separation was between WT on the one hand and both mutants on the other hand (Fig 6A). Cell types in the WT separate partially, with the A and E types being clearly identifiable, while the other types are mixed together. For the mutants, only the A cells build a clear cluster. None of the mutant samples groups with the WT cell types. We deepened the analysis by using the three types A, E and R of the WT as references to examine the WT I and B, and to compare pairs of genotypes WT *vs etl* on the one hand, and WT *vs kna* on the other hand. We first identified genes for which expression was the most variable between A, E and R types in the WT. This was done by computing the Variance Ratio (VR), i.e. dividing the variance between types by the variance between replicates. The 2000 genes having the highest VR (> 5) were retained. As expected, these genes allowed to separate the three WT cell types by PCA, and clustered the replicates for each type. Using the coefficients computed from this first step, we plotted the other cell types of the WT (Fig 6B). Both cell types I and B located in the centre of the PCA space. This fits our expectation because I is a transitory stage corresponding to cells in the process of rounding, and B resumes apical growth through branching. To find out which parameter is paramount in the control of cell differentiation, we performed PCA with the WT and *etl* on the one hand, and with WT and *kna* on the other hand. Fig 6C shows that the 3 spatial domains in *etl* clustered with the R cells of the WT. This suggests that cell shape controls cell fate, as all spatial domains were made of round cells in *etl*. Cell age might count too, as *etl* cells in the A and S spatial domains are all more mature than A and E cells from the WT.

**Figure 6:**
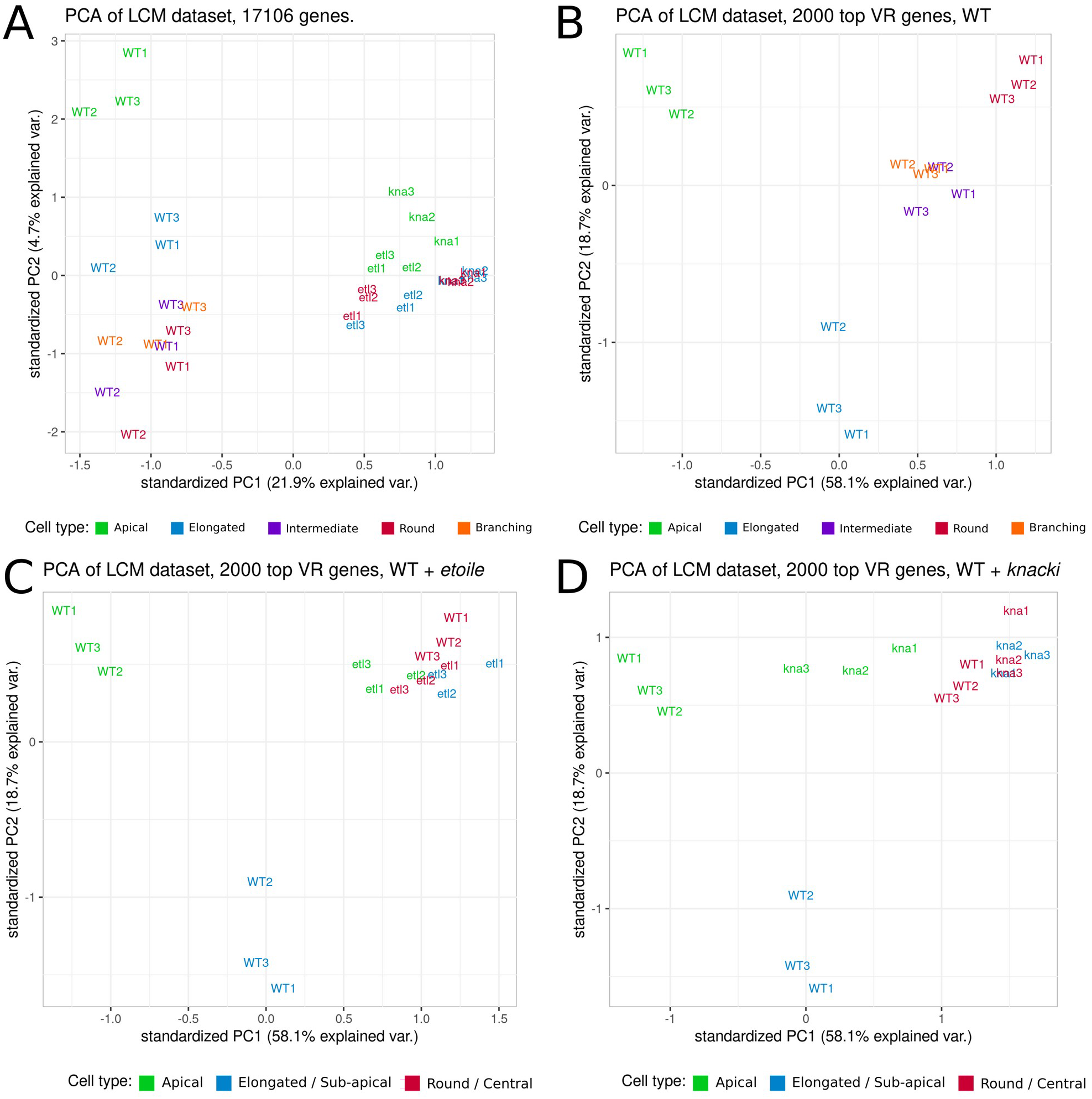
Mutants lost cell-type specific transcriptome. Principal Component analysis (PCA) of gene expression was used to compare cell domain expression patterns. (A) All genes and all samples were included in the analysis. (B-D) Only 2000 genes, identified as the most discriminant ones for A, E, R in WT, were considered in the analysis, and the principal components, computed only on these three types, were kept for further plotting. (B) adding I and R types, to compare the five WT cell types. (C) adding the three *etoile* spatial domains. (D) adding the three *knacki* spatial domains.

However, PCA including WT and *kna* led to a strikingly different conclusion: none of the *kna* spatial domain clustered with elongated cell types from the WT (Fig 6D). Instead, *kna* S and *kna* C domains clustered with WT R cells. Only the *kna* A spatial domain displayed some vicinity with the WT A cells, when compared with the other two domains. Therefore, PCA analysis with *kna* refutes the hypothesis that cell shape alone controls cell fate, at least when the transcriptome is used as a reflection of cell identity.

This PCA confirmed that *etoile* and *knacki* are more similar with each other than any of these mutants with the WT (*cf* Fig 5). Therefore, summing the PCA analysis of both mutants, neither the spatial location within the tissue nor the cell shape control gene expression in these mutants. Cell maturity (age) could be a factor for *etl*, but this does not apply to *kna*.

To get more answers, we analysed the GO-term differentially represented in these mutants. We found only two GO-terms, related to translation activity (GO:0003735: “Structural constituent of ribosome” and GO:0006412: “translation”) that were > 100 times over-represented in the pool of genes that were up-regulated in WT A cells compared to *kna* A domain (P-value < 5×10^−3^; Suppl File 2). This is very surprising, because WT A cells are amongst the cells that do not over-expressed genes belonging to this GO-term. The fact that *kna* A cell does it even less than its WT counterpart is puzzling.

## Discussion

By analysing the transcriptome of cells along the WT filament, we showed that transcriptome activity followed changes in cell identity, which involves simultaneous transition of three parameters: age, location and shape (Fig 2B). The two morphological mutants *etl* and *kna* display opposite cell shapes – bulky to spherical in *etl* and homogeneously elongated and cylindrical in *kna–* associated with different combinations of ages and positions along the filament. Therefore, comparative analyses of transcriptomes of the WT cell types, and specific spatial domains of the two mutants offer the opportunity to split up the contribution of the three parameters. The comparison of *kna* central domain to WT R cells and *etl* central domain (Fig 5E) is of particular interest, because it is the only case where pairs of types/domains differ by exactly one parameter, namely the shape (Fig 2A). The fact that only one gene is DE between WT and *kna*, and no gene at all is DE between *kna* and *etl* constitutes a clear indication that there is no transcriptome signature attached to shape. Therefore, other parameters must be overlapping the major effect of cell shape modification on the control of gene regulation.

Additional factors like the cell size could be candidate. Cell size was shown to control gene expression in the zygotic genome activation in metazoan blastomeres (Chen et al., 2020). In *Ectocarpus*, the volume of R cells before cell division can be up to 3 times higher than that of E cells (it is 27% higher in R cell after cell division) and therefore, this difference could maybe result in a change in the genetic program through e.g. a modification of 3D genome organisation (Aboelnour and Bonev, 2021). However, while their transcriptome signature is close to the WT R cells, *kna* cells have a similar volume as WT E cells. Therefore, the difference in volume does not seem to be a key parameter in the control of gene expression pattern, at least in a mutant context.

Noticeably, the two mutants lose the E cell type identity. In the PCA analysis, the S cell domainn of both *etl* and *kna* clustered with the WT R cells. This is comprehensible for *etl*, that has no elongated cells, but because of the phenotype of *kn*a, we expected its S spatial domain to cluster with WT E. In fact, when performed with all available genes, PCA first separated the cell types of the two mutants from those of the WT. When only the genes that efficiently sorted the three WT typical cell types were used, all mutant spatial domains had an expression pattern close to the WT R cells. Hence, it could turn out that this study actually tells us more about the nature of the WT R cells than on the factors controlling gene expression. A challenging hypothesis would then be that the R cell stands for a default status in the WT *Ectocarpus*. Progressive and time-dependent cell rounding from cylindrical and very polarised apical cells could be viewed as a relaxing process where cells release constraints underlying their initial anisotropic shape. In this hypothesis, *kna* cells would remain physically “locked” in a constrained state, while *etl* cells would appear insensitive to the putative constraining factor, but both would be in this genetic default status because they are mutants. Interestingly, work in bacteria, in which mechanisms leading to the formation of the rod-shaped versus coccal cell shapes, were compared (Pinho et al., 2013) fuels this hypothesis. While rod-shaped cells might be considered intuitively as having evolved from cocci genera, phylogenetic studies actually support the opposite: coccus morphology is more recent and evolved from rod-shaped genera. In eukaryotes, cell rounding is a transitory stage allowing the symmetrical distribution of cellular components during mitosis before cells recover specific cell shapes (Godard et al., 2020; Lancaster et al., 2013; Stewart et al., 2011). Furthermore, using non-linear poro-elastic models, we showed that cell rounding in *Ectocarpus* can easily be accounted for if the cell is turgid (Jia et al., 2017), which it is (Rabillé et al., 2019). Therefore, it is conceivable that cell rounding in *Ectocarpus* corresponds to a release of the cell wall mechanical constraints that are actively set up in the apical cell. These constraints would be impaired in both mutants, turning all mutant cells into a default status. This situation echoes the impact of cell length on the transcriptome of cardiomyocytes, in which both shortening or lengthening of cells results in a similar reduction in expression of signaling-related genes (Haftbaradaran Esfahani et al., 2020). However, it should be noted that in *Ectocarpus*, the cells of the mutants differ from the WT R type in several DE genes, giving them their own signature. Altogether, it appears difficult to disentangle relationships between cell shape, cell age and cell position, and morphological mutants might be too drastic an approach in *Ectocarpus* to address this question. Approaches with mutants clearly impaired in differentiation timing, or showing abnormal positioning of each different cell type might prove fruitful, but difficult to obtain.

Is this a particular case? Among the other filamentous organisms, mosses are the ones with the highest morphological resemblance to *Ectocarpus*. Adult *Physcomitrium* (former *Physcomitrella*) gametophytes are comprised of branched uniseriate filaments growing apically. Interestingly, the first cell types differentiating from the spore, the chloronemata, are slow growing and committed mainly to photosynthetic assimilatory function, while the second cell type, the caulonemata, are fast growing and forage for water and nutrients (Coudert et al., 2019; Cove, 2005). Nicely, this spatial distribution of functions is reminiscent of what is observed in *Ectocarpus*. In both cases, photosynthetic, energy producing cells are located in the centre of the filaments, while actively growing and foraging cells are in the extremities. However, while cell differentiation is centripetal in *Ectocarpus*, it is centrifugal in *Physcomitrium* (note that reverse differentiation from caulonemata to chloronemata occurs later in development (Cove et al., 2006). Auxin controls the caulonemata differentiation through PIN-mediated polarized transport towards the apex and a similarly orientated gradient of auxin has been displayed in *Ectocarpus* sporophytic filaments (Le Bail et al., 2010). Xiao et al. (2011) analysed gene expression in the gametophytes of the moss *Physcomitrium patens*, through RNA-seq and DEG analyses on a temporal series of gametophytic filaments. They found many genes involved in metabolic pathways and energy metabolism, in addition to several transcription factors. Their study was not at the single cell type level, but encompassed the whole filaments taken at different developmental stages, in addition to laser capture microdissected caulonemal and chloronemal cells, in which they found ∼ 400 genes DE between the caulonemata and the chloronemata (half up-regulated). This number is close to the number of DE genes that we found between the R and the E cells in WT *Ectocarpus*. In contrast to their study, we did not identify any GO-term specific for transcription factors. The micromanipulation based scRNA seq technique developed by Kubo et al. (2019) on *Physcomitrium* provides an opportunity to obtain transcriptomics data on each individual cell of the protonemata, and extend the comparison with *Ectocarpus* cell differentiation. In conclusion, *Ectocarpus* allowed us to examine the fundamental process of cell differentiation in the Stramenopile lineage for the first time with such a high spatial resolution. Additional cell differentiation mutants might be necessary to refine our current understanding of the underlying mechanisms of cell fate in this alga.

## Supporting information

Supplementary File 1 - DE genes

Supplementary File 2 - GO-terms

Supplementary movie 1 - Development

## Acknowledgment

PEPS CNRS (B.C.) and ERC (J.A.L.) funded the transcriptomic sequencing. D.S.M. was funded by an ERC AdG (EDIP) grant to J.A.L.

The authors are thankful to Elodie Rolland for the maintenance of the algal cultures.

## Author contribution

B.B. performed the sequence analyses, C.D. and B.B. performed the time-lapse monitoring of cell rounding. C.S. performed the Q-PCR. B.C. and D.S.M. performed laser capture experiments. D.S.M. performed the RNA extraction and amplification. B.C. designed the biological experiments and wrote the initial version of the paper. All authors contributed to the final version of the manuscript.

## Supplementary data

### Supplementary Materials & Methods

#### Quantitative-RT-PCR

Method: Each WT cDNA sample was amplified using 5’ and 3’ oligonucleotides (Eurogentec) specific of two distinct exons for each gene. Oligonucleotides were designed with the software Perlprimer (Marshall, 2004). Genes were Armadillo-like helical (Ec-12_007670), long chain acyl-coA synthetase (Ec-12_008720), Acyl-CoA dehydrogenase (Ec-14_001600), conserved unknown protein (Ec-14_001610), expressed unknown protein (Ec-17_004270), Mannuronan C-5-epimerase (Ec-19_003150), UDP-glucose/GDP-mannose dehydrogenase, N-terminal (Ec-19_004990), Catalase (Ec-26_000310), Secreted protein similar to EsV-1-163 (Ec-26_003200), SGNH hydrolase-type esterase domain (Ec-26_004720), Carbohydrate-binding WSC (Ec-28_003030). List of oligonucleotides is given in Suppl Table 2. cDNA amplification took place in 5µL composed of 2.5 µL SYBR green I master (Roche), 0.2 µL of each oligonucleotide (10 µM initial concentration), and 0.6 ng/µL cDNA (2.1 µL). Amplification programme followed by the LightCycler 480 multiwell plate 384 (Roche) was Preincubation – 1 cycle: 95°C for 5min; Amplification for 45 cycles with 10sec at 95°C, 30sec at 60°C, 15sec at 72°C; Melting curves for 1 cycle 5min at 95°C, then 1min at 65°C and finally progressive heating up to 97°C with a fluorescent recording every second. Sample was then cooled down to 40°C and maintained for 30sec. Absolute transcript copy number was calculated by comparison with a concentration dilution series made of *Ectocarpus* sp. genomic DNA. Five concentration points, corresponding to 22376 copies (5.25 ng), 8951 copies (1.05 ng), 895 copies (0.105 ng), 89 copies (0.0105 ng) and 9 copies (0.00105 ng). The absolute number of transcripts quantified in the cDNA samples was normalised to the geometric average of the three best reference genes identified in Le Bail et al. (2008). The log2 of this value was averaged over the 3 biological replicates.

### Supplementary results

#### Abundance of reads

Initial counting of the raw data showed that the five cell types of the WT filaments contain comparable number of reads, with the mutants *etoile (etl)* and *knacki (kna)* having ∼1.5 times more reads, except for R cells in *kna* (written thereafter ‘knaR’) (Suppl Fig 1). Similar abundance between samples was maintained after RNA trimming that removed the low quality sequences (Suppl Fig 2). As RNA amplification involved both polyA and random priming, many non-conding RNAs, especially ribosomal RNAs were present in the initial dataset and were removed at this step. Representativity of the remaining non-rRNA reads in each sample ranged from 2.9 to 12.2% (average 6.6%) of the total read numbers per sample (Suppl Fig 3). These non-rRNA reads were more abundant in mutant samples compared to WT samples (except in knaR), probably because the total number of reads in these samples was higher (Suppl Fig 4). Despite this discrepancy, the ratio between the most and the less abundant over the whole dataset was less than 2, what we consider suitable for subsequent analyses. When the replicates of the same sample type were pooled, the number of exploitable reads ranged between 1,767,722 and 4,866,42 per cell type, with an average of ∼ 3,370,000 non-rRNAs (Suppl Fig 5; Suppl Table 1). Therefore, we considered that the homogeneity of our dataset was high enough to proceed the analysis further. In a third step, these RNAs were mapped onto the *Ectocarpus sp*. genome and transcriptome (Cormier et al., 2017). 86.3% of the non-rRNAs identified above mapped on the *Ectocarpus* genomic sequence. They were localised in predicted CDS (40%), predicted intergenic regions (30%) and introns (20%). Therefore, ultimately ∼ 4.5% of the total initial read number mapped predicted CDS, either on the nuclear (13.976 millions reads), chloroplastic (12.779 millions reads) or mitochondrial CDS (10.376 millions reads).

#### QT-RT-PCR

We used Q-RT-PCR amplification to check the reliability of the NGS data on a series of genes found to be significantly differentially expressed between the A and R cell types. Five genes showing a strong expression bias from the NGS analysis (fold change > 4 with a FDR < 10^−13^) for A cell, namely Mannuronan C-5-epimerase (Ec-19_003150), UDP-glucose/ GDP-mannose dehydrogenase, N-terminal (Ec-19_004990), Catalase (Ec-26_000310), Secreted protein similar to EsV-1-163 (Ec-26_003200), Carbohydrate-binding WSC (Ec-28_003030) were chosen. Similarly 6 genes showing a significant expression bias for R cells from the NGS analysis, namely Armadillo-like helical (Ec-12_007670), long chain acyl-coA synthetase (Ec-12_008720), Acyl-CoA dehydrogenase (Ec-14_001600), conserved unknown protein (Ec-14_001610), expressed unknown protein (Ec-17_004270), SGNH hydrolase-type esterase domain (Ec-26_004720) were chosen. The absolute number of transcripts for these 11 genes was quantified by Q-RT-PCR and was normalised with 3 reference genes. Two exons were amplified for each gene, and the ratio of expression between the R cells and the A cells was calculated for each exon. It was compared to the same ratio calculated from the number of reads obtained from NGS. Suppl Table 4 and Suppl Fig 6 show that for each exon tested, the sign of log(fold-change) was the same with Q-RT-PCR than with NGS, meaning that under/over expression is found to vary similarly. Nevertheless, a student t-test failed to show evidence for a significant difference (α = 5×10^−2^) between the values obtained and 0, for 16 out of 21 assays (Suppl Table 4). This was obviously due to a high variability between replicates. Conversely, all but two exons displayed a non-significant difference with the NGS result (Suppl Table 4). In conclusion, the results of this analysis constitute a qualitative confirmation of the differential expression pattern found by NGS analysis.

### Supplementary figures

**Supplementary figure 1:**
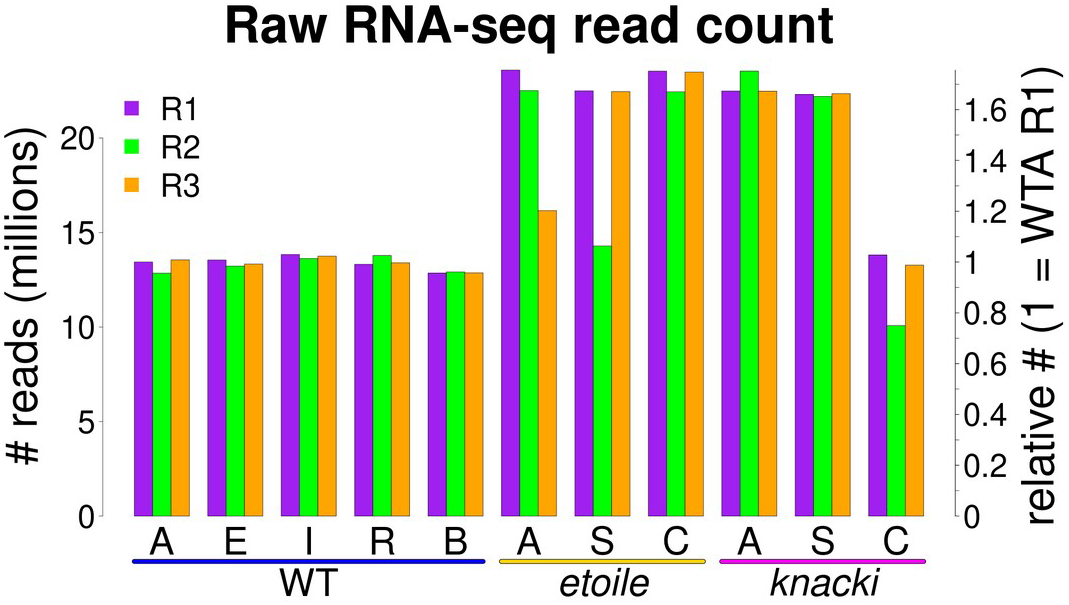
Number of reads in each sample, before trimming. R1, R2, R3 represent the three independent replicates. Right scale: normalised to the first replicate (R1) of the WT apical cell transcriptome. *Etoile and knacki* mutant samples contained in average more reads than the WT samples (exception for *knacki* C).

**Supplementary figure 2:**
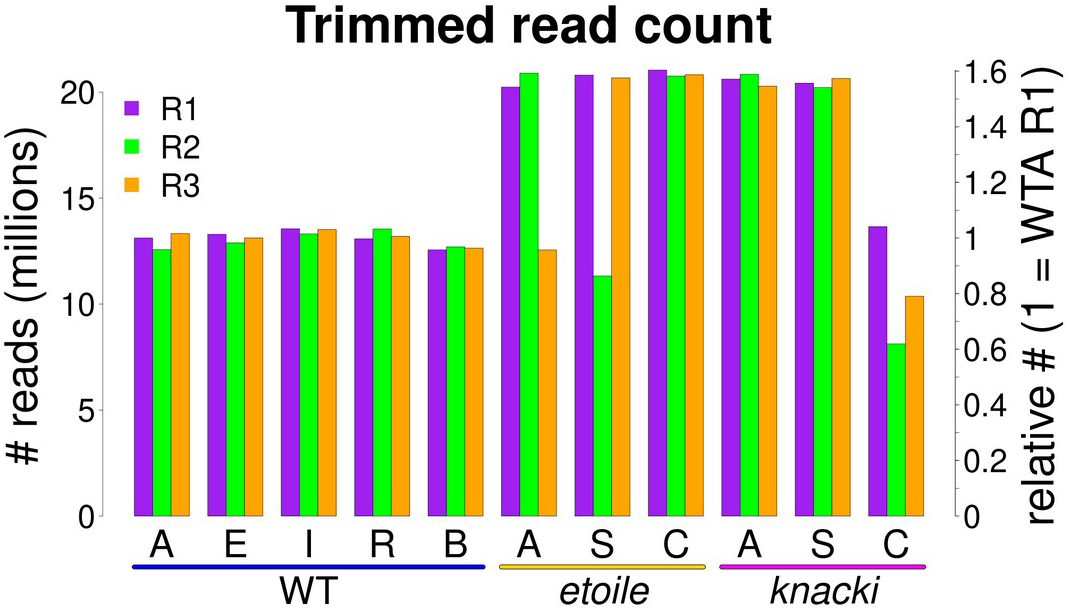
Abundance after RNA trimming in all the samples. Right scale: normalised to the sample corresponding to the WT Apical cells, R1.

**Supplementary figure 3:**
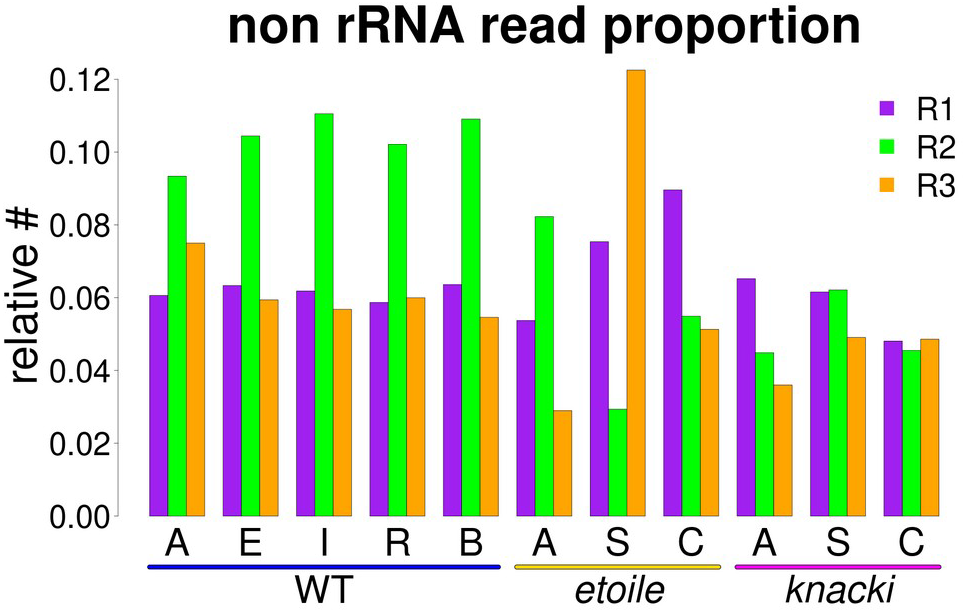
Representativity of non rRNAs in the total reads dataset. Ratio ranges between 3 and 12 %. *etoile* A and *etoile* S were the most disparate, and displayed the lowest % representativity in one of their 3 replicates (*etoile* A, R3 and *etoile* S, R2, respectively). in WT, R2 showed systematically a higher representativity in all cell types. These discrepancy might be due to the RNA amplification step.

**Supplementary figure 4:**
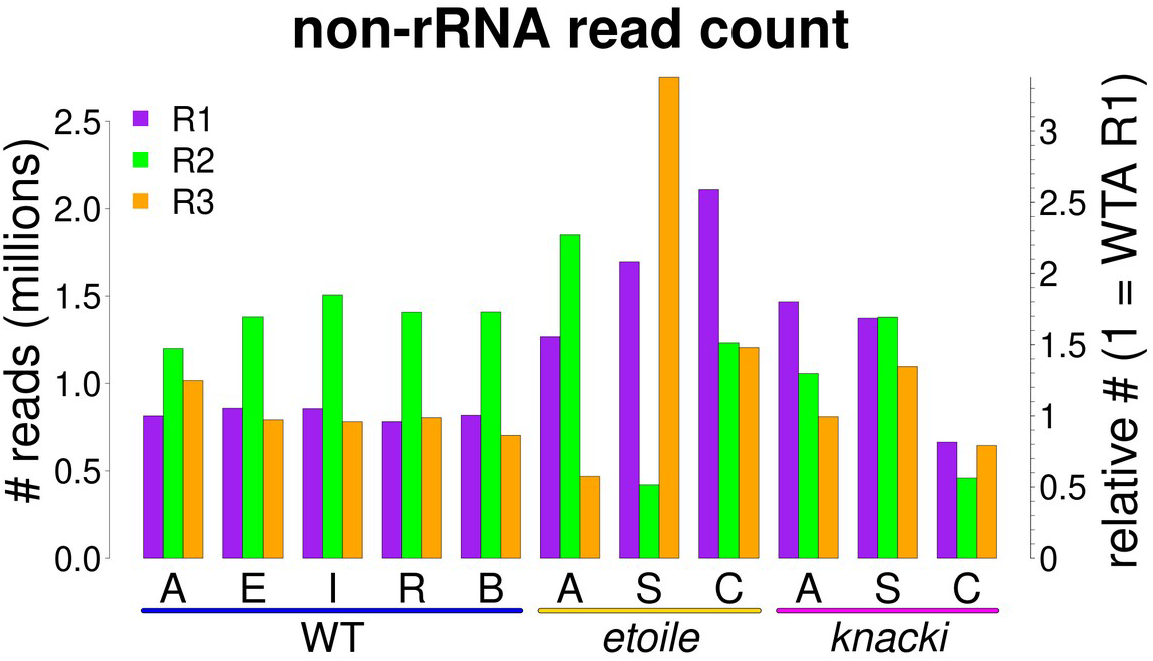
Number of non rRNA reads per sample. Right scale normalised to WT A, R1.

**Supplementary figure 5:**
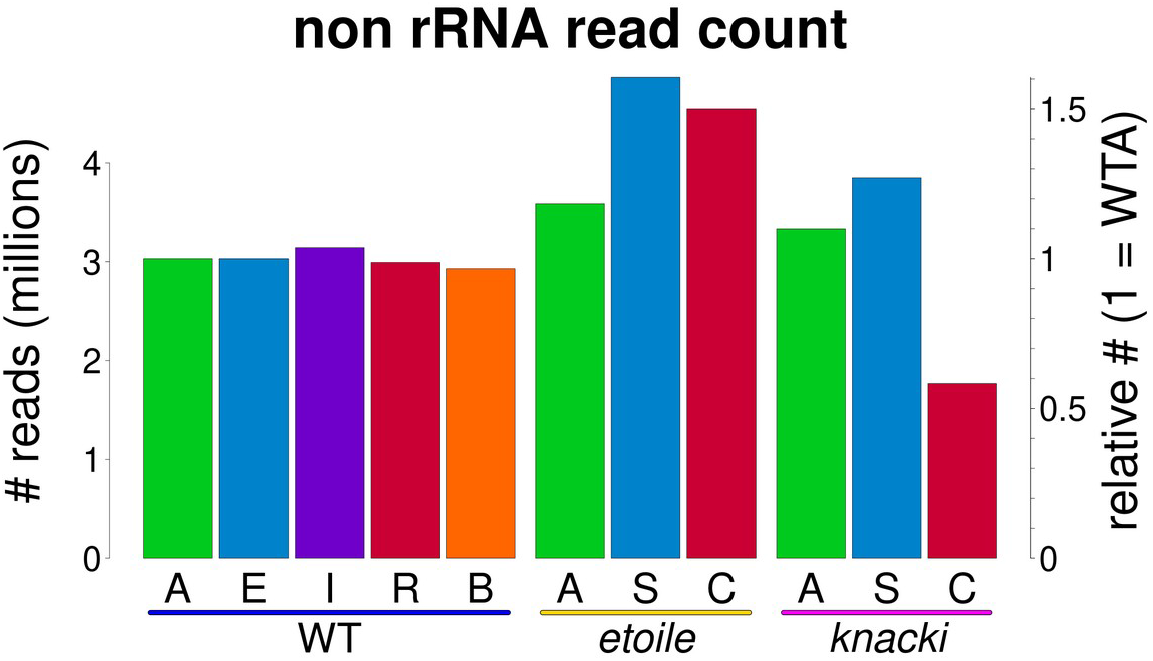
Total number of non-rRNA reads per cell type. Right scale: normalised to the WT A amount. In average, the highest amount was available in the mutant *etoile*.

**Supplementary figure 6:**
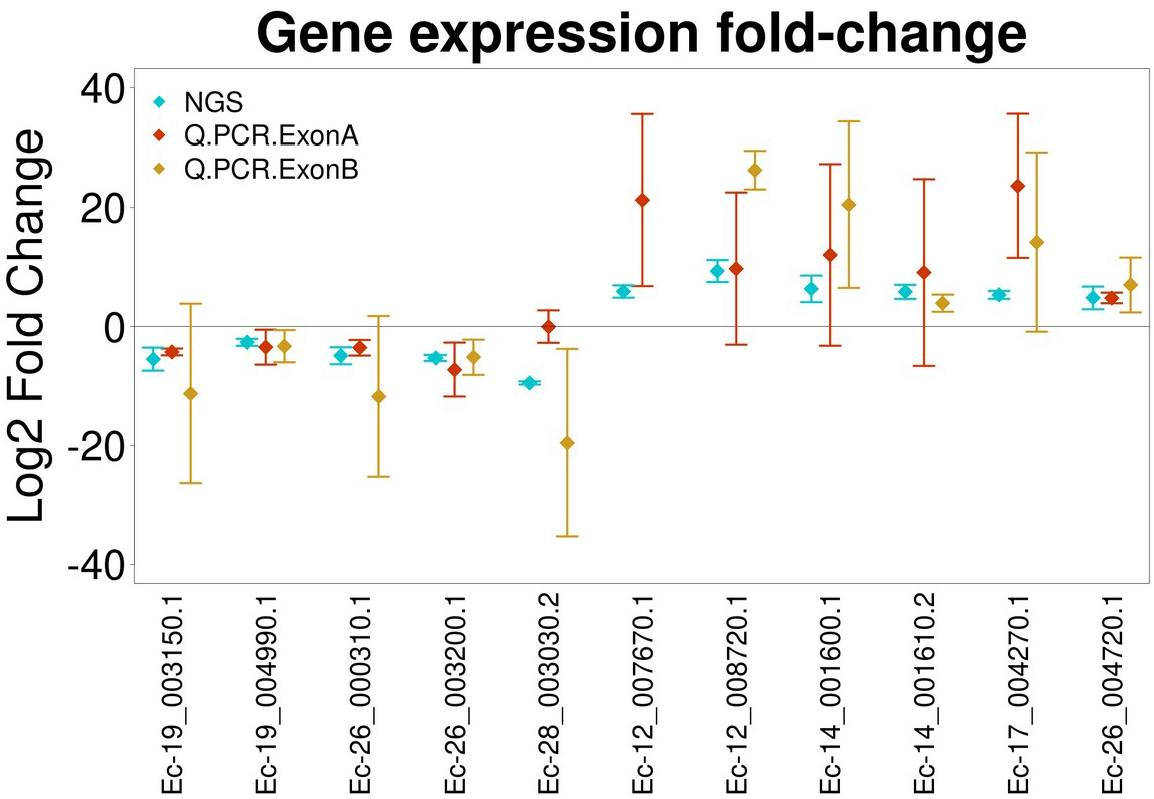
Transcript level comparison between NGS and Q-PCR quantification for 11 genes in the two WT cell types A and R. For the Q-PCR, two exons were amplified for each gene when possible. *Ectocarpus* genes exon are 100bp in average (Cock et al., 2010) and for gene Ec-12_007670.1 a second exon could not be amplified with an PCR efficiency higher than 90%, as requested to ensure quantification.

**Supplementary figure 7:**
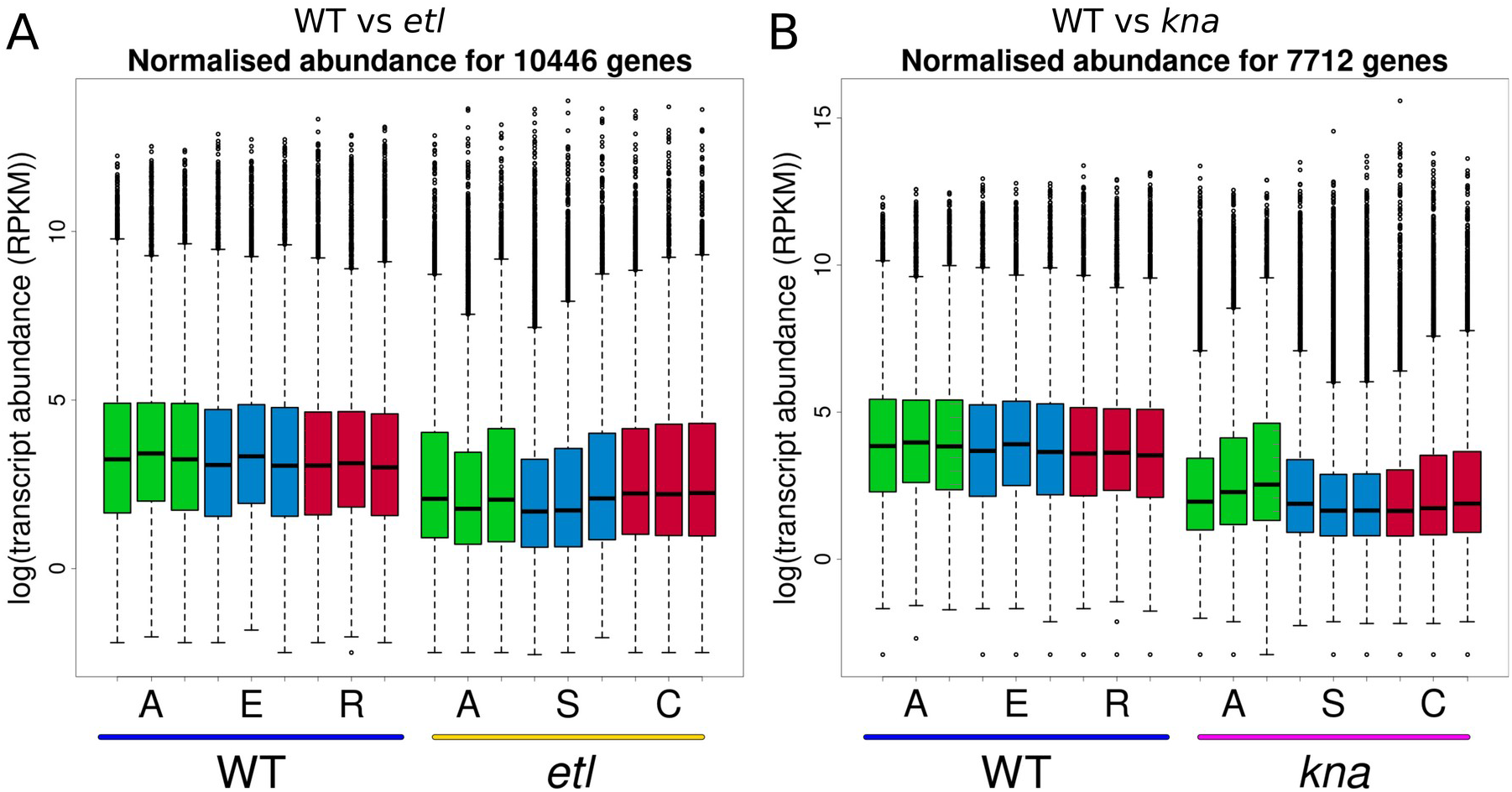
Comparison of transcript abundance in the 3 replicates of the 3 cell types or spatial domains, expressed in log2 of the RPKM, between (A) WT and *etl* (10446 genes in common), and (B) WT and *kna* (7712 shared genes).

### Supplementary tables

**Supplementary table 1:** Raw number of RNA-seq reads for all samples in the analysis.

| Strain | Type | Repl 1 | Repl 2 | Repl 3 | Sum |
| --- | --- | --- | --- | --- | --- |
| WT | A | 13118492 | 12566817 | 13332084 | 39017393 |
| WT | E | 13292636 | 12887627 | 13122394 | 39302657 |
| WT | I | 13553607 | 13312260 | 13518700 | 40384567 |
| WT | R | 13080925 | 13541901 | 13193853 | 39816679 |
| WT | B | 12559509 | 12700885 | 12638891 | 37899285 |
| *etl* | A | 23590504 | 22502562 | 16159782 | 62252848 |
| *etl* | S | 22493142 | 14286129 | 22457810 | 59237081 |
| *etl* | C | 23537604 | 22440294 | 23483018 | 69460916 |
| *kna* | A | 22485257 | 23537888 | 22477271 | 68500416 |
| *kna* | S | 22307896 | 22197196 | 22346555 | 66851647 |
| *kna* | C | 13812781 | 10077094 | 13276106 | 37165981 |

**Supplementary table 2:**
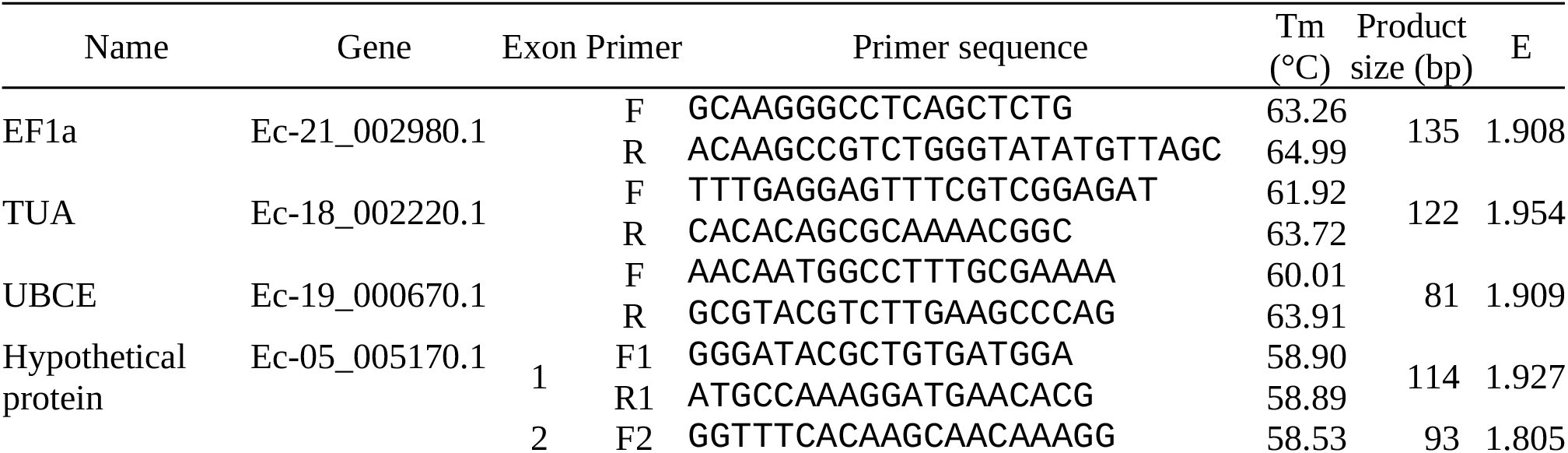
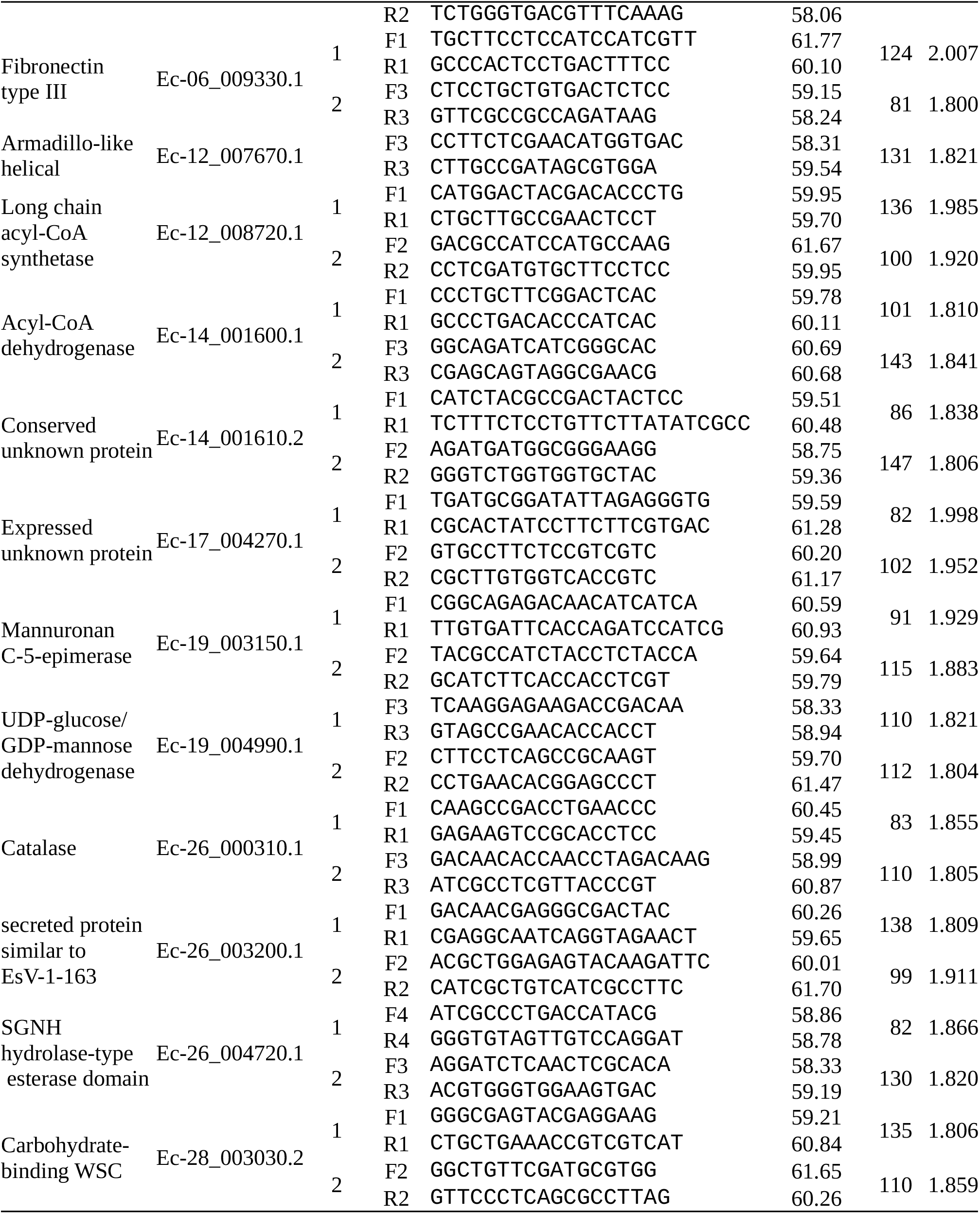
List of oligonucleotides used for RT-Q-PCR. E: PCR efficiency.

**Supplementary table 3:** Sporophytic- and gametophytic-biased and specific genes expressed in the early stage sporophyte. GS: Gametophyte-specific genes; GA: gametophyte-biased genes; UB: ubiquitously expressed genes; SP: Sporophyte-biased genes; SS: Sporophyte-specific genes. --: genes absent from the study of Lipinska et al. (2017). A, E, I, R, B are the letters given to each cell types identified in the WT *Ectocarpus*. Number of genes of each type, as defined from Lipinska et al (2017), are indicated on the central column of the table (Lip), while percentage of their representativity within them is indicated in the right-hand side.

|  | # genes expressed |  |  |  |  | Lip | % of Lip genes expressed |  |  |  |  |
| --- | --- | --- | --- | --- | --- | --- | --- | --- | --- | --- | --- |
|  | A | E | I | R | B |  | A | E | I | R | B |
| GS | 32 | 30 | 37 | 32 | 24 | 250 | 0.00 | 12.00 | 14.80 | 12.80 | 9.60 |
| GA | 1771 | 1679 | 1800 | 1706 | 1456 | 3855 | 45.94 | 43.55 | 46.69 | 44.25 | 37.77 |
| UB | 8145 | 8102 | 8245 | 8053 | 7464 | 11232 | 75.52 | 72.13 | 73.41 | 71.70 | 66.45 |
| SP | 1663 | 1646 | 1706 | 1684 | 1567 | 2001 | 83.11 | 82.26 | 85.26 | 84.16 | 78.31 |
| SS | 45 | 47 | 52 | 52 | 42 | 96 | 46.88 | 48.96 | 54.17 | 54.17 | 43.75 |
| – | 523 | 483 | 522 | 496 | 439 |  |  |  |  |  |  |

**Supplementary table 4:** Comparison between expression variations obtained by NGS and S-PCR. p-values <0.05 are denoted in bold.

| Gene | Exon | NGS | Q-PCR |  |  |  | p-value for t-test QPCR vs... |  |
| --- | --- | --- | --- | --- | --- | --- | --- | --- |
|  |  |  | R1 | R2 | R3 | Mean | 0 | NGS |
| Ec-19_003150.1 | A | -4.59 | -4.14 | -3.79 | -4.89 | -4.27 | **0.006** | 0.432 |
|  | B | -4.59 | -28.38 | -5.36 | 0.00 | -11.25 | 0.326 | 0.524 |
| Ec-19_004990.1 | A | -2.20 | -6.77 | -2.50 | -1.13 | -3.47 | 0.178 | 0.534 |
|  | B | -2.20 | -6.39 | -2.23 | -1.30 | -3.31 | 0.169 | 0.553 |
| Ec-26_000310.1 | A | -4.22 | -5.08 | -2.66 | -2.99 | -3.58 | **0.042** | 0.485 |
|  | B | -4.22 | -6.04 | -2.03 | -27.18 | -11.75 | 0.271 | 0.436 |
| Ec-26_003200.1 | A | -4.82 | -12.46 | -4.61 | -4.62 | -7.23 | 0.110 | 0.454 |
|  | B | -4.82 | -8.45 | -4.38 | -2.69 | -5.17 | 0.094 | 0.855 |
| Ec-28_003030.2 | A | -8.96 | -3.17 | 1.54 | 1.59 | -0.01 | 0.994 | **0.030** |
|  | B | -8.96 | -30.57 | -1.48 | -26.54 | -19.53 | 0.165 | 0.365 |
| Ec-12_007670.1 | A | 5.93 | 25.82 | 5.05 | 32.70 | 21.19 | 0.126 | 0.208 |
| Ec-12_008720.1 | A | 9.92 | 0.00 | 4.97 | 24.20 | 9.72 | 0.318 | 0.981 |
|  | B | 9.92 | 22.52 | 27.85 | 28.34 | 26.24 | **0.005** | **0.013** |
| Ec-14_001600.1 | A | 5.68 | 2.29 | 4.16 | 29.57 | 12.01 | 0.306 | 0.547 |
|  | B | 5.68 | 29.56 | 4.38 | 27.35 | 20.43 | 0.126 | 0.208 |
| Ec-14_001610.2 | A | 5.74 | 0.00 | 0.00 | 27.17 | 9.06 | 0.423 | 0.749 |
|  | B | 5.74 | 5.56 | 3.30 | 2.89 | 3.92 | **0.042** | 0.159 |
| Ec-17_004270.1 | A | 5.70 | 30.05 | 9.68 | 31.03 | 23.59 | 0.077 | 0.124 |
|  | B | 5.70 | 4.28 | 6.74 | 31.46 | 14.16 | 0.244 | 0.432 |
| Ec-26_004270.1 | A | 4.85 | 5.36 | 5.26 | 3.78 | 4.80 | **0.011** | 0.931 |
|  | B | 4.85 | 4.69 | 3.95 | 12.27 | 6.97 | 0.120 | 0.509 |

**Supplementary table 5:** Number of differentially expressed (DE) genes for pairwise comparisons between cell types / domains, for all cases where differences occur for one or two parameters among (E) elongation, (A) age and (P) position.

| Cell 1 | vs Cell2 | Parameter(s) | # DE genes |
| --- | --- | --- | --- |
| WT R | *kna* C | E | 1 |
| *etl* C | *kna* C | E | 0 |
| WT A | *etl* A | E + A | 137 |
| WT E | *etl* S | E + A | 19 |
| *etl* A | *kna* A | E + A | 3 |
| *etl* S | *kna* S | E + A | 2 |
| WT E | *etl* A | E + P | 125 |
| WT I | *etl* S | E + P | 7 |
| *kna* S | *etl* A | E + P | 2 |
| WT I | *etl* A | P + A | 156 |
| WT R | *etl* S | P + A | 11 |
| WT E | *kna* C | P + A | 2 |

### Supplementary files

<u>Supplementary file 1:</u> DEGenes_Ectocarpus.ods shows differentially expressed genes between cell types in WT, and in each mutant *etoile* and *knacki*.

<u>Supplementary file 2:</u> GOterms_Ectocarpus.ods shows GO-terms having enrichment in lists of differentially expressed genes between cell types in WT, and in each mutant *etoile* and *knacki*.

## Supplementary movie

**Supplementary movie 1:**
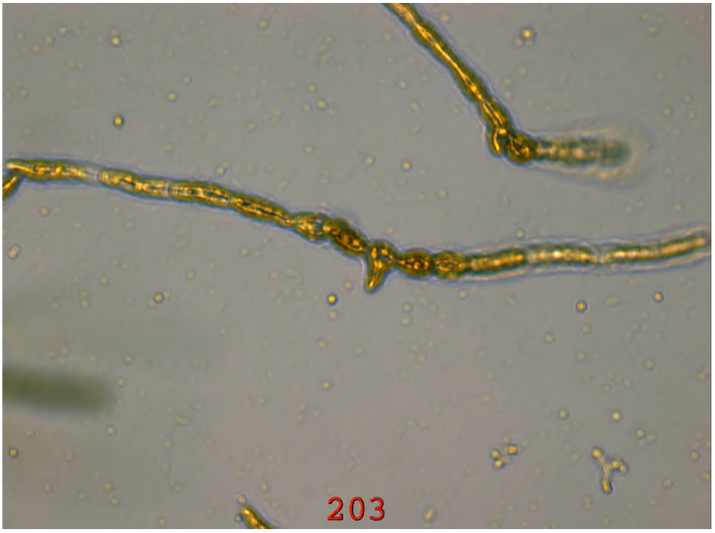
EctocarpusDevelopment_01.webm shows a time-lapse movie of a young WT *Ectocarpus sp*. sporophyte. Apical (A) cells grow and periodically divide to give new sub-apical Elongated (E) cells. These E cells then get rounder, passing through an Intermediate (I) stage before they eventually become Round (R). Occasionally, cells develop lateral branches: they are called Branched (B) as they give rise to a novel filament led by an A cell. Time is given in hours.

## Notes

### Competing Interest Statement

The authors have declared no competing interest.

